# Quantifying spontaneous metastasis in a syngeneic mouse melanoma model using real time PCR

**DOI:** 10.1101/136945

**Authors:** Wentao Deng, Sarah L. McLaughlin, David J. Klinke

## Abstract

Modeling metastasis in vivo with animals is a priority for both revealing mechanisms of tumor dissemination and developing therapeutic methods. While conventional intravenous injection of tumor cells provides an efficient and consistent system for studying tumor cell extravasation and colonization, studying spontaneous metastasis derived from orthotopic tumor sites has the advantage of modeling more aspects of the metastatic cascade, but is challenging as it is difficult to detect small numbers of metastatic cells. In this work, we developed an approach for quantifying spontaneous metastasis in the syngeneic mouse B16 system using real time PCR. We first transduced B16 cells with lentivirus expressing firefly luciferase *Luc2* gene for bioluminescence imaging. Next, we developed a real time quantitative PCR (qPCR) method for the detection of luciferase-expressing, metastatic tumor cells in mouse lungs and other organs. To illustrate the approach, we quantified lung metastasis in both spontaneous and experimental scenarios using B16F0 and B16F10 cells in C57BL/6Ncrl and NOD-Scid Gamma (NSG) mice. We tracked B16 melanoma metastasis with both bioluminescence imaging and qPCR, which were found to be self-consistent. Using this assay, we can quantitatively detect one Luc2 positive tumor cells out of 10^4^ tissue cells, which corresponds to a metastatic burden of 1.8x10^4^ metastatic cells per whole mouse lung. More importantly, the qPCR method was at least a factor of 10 more sensitive in detecting metastatic cell dissemination and should be combined with bioluminescence imaging as a high-resolution, end-point method for final metastatic cell quantitation. Given the rapid growth of primary tumors in many mouse models, assays with improved sensitivity can provide better insight into biological mechanisms that underpin tumor metastasis.

## Introduction

Cancer metastasis, which is the migration of malignant cells from the primary site of origin to distant tissues, is the primary cause of death among cancer patients, responsible for as much as 90% of cancer-associated mortality ^1,2^. For instance in patients with melanoma, the 5-year survival rate drops from 98% for localized melanoma to 63% for regional and 17% for distant stage melanoma ^3^. While genetic alterations are critical for malignant transformation, identifying how specific genetic alterations interact with microenvironmental signals to enable metastasis to distant vital organs remains a challenge.

Tumor metastasis is a multistep cascade that starts with local invasion into the surrounding tissue and intravasation into nearby blood and lymphatic vessels ^4,5^. The tumor cells then translocate to distant tissues, exit from the bloodstream (extravasation), and interact with a new tissue microenvironment forming micrometastases that eventually grow into macroscopic tumors (colonization) ^4,5^. The biological complexity that characterizes metastasis requires a thorough understanding of each step, and modeling metastasis in vivo with animals will provide critical insight for both the mechanism and treatment.

Compared to genetically engineered mouse models that exhibit variable phenotypes and require prolonged periods before metastases may appear, transplantable mouse models, including both syngeneic and xenograft models, are widely used to recapitulate the entire tumor metastatic process ^6,7^. Depending on the study design, transplantation models assay either spontaneous metastasis or experimental metastasis based on how the tumor cells are delivered to the recipient animals. Spontaneous metastasis assays need to establish a primary tumor and allow it to grow and metastasize, whereas experimental metastasis assays circumvent the initial growth, invasion and intravasation steps by directly injecting tumor cells into the circulation. Experimental metastasis assays are fast, reproducible and consistent, but spontaneous metastasis assays provide an opportunity to study the whole metastatic cascade and many aspects that are bypassed using experimental metastasis models ^6,7^. However, spontaneous metastasis assays are less commonly used due to low tumor metastatic rate and difficulty detecting for the presence of metastatic tumor cells. Although orthotopic tumor transplantation or removal of the primary tumor could promote the metastatic phenotype, the traditional methods to assess tumor metastasis, such as morphometric quantitation of lung colonies (nodules), are not very sensitive, accurate, nor quantitative, especially when metastatic tumor cells cannot produce macroscopically visible colonies in secondary organs.

Recent technology development to track and quantify tumor cells provides more sensitive methods for detecting micrometastases in spontaneous metastasis assays. One set of noninvasive imaging methods take advantage of genetically introduced imaging reporters in cancer cells to track their location in vivo. These techniques include fluorescence, bioluminescence, positron emission tomography (PET), and single-photon emission computed tomography (SPECT) ^8–10^. Among these, bioluminescence imaging is the most commonly used system, which uses genetically introduced luciferase to catalyze a light-producing reaction from injected substrate. Bioluminescence imaging provides a relatively simple, robust, cost-effective, and sensitive method to monitor the biological processes in vivo, but detection resolution and outcomes depend on substrate delivery and substrate pharmacokinetics, in addition to tumor location, tumor cell viability, and penetration of light through animal tissues ^8,10^. As a result, tumor burden derived from the strength of the bioluminescence signal is only a semi-quantitative measurement ^8,10^.

Another set of new methods for metastatic tumor cell detection are the real time quantitative PCR (qPCR) based assays that use species-specific oligomer primers or probes to quantify species-specific genomic DNA or cDNA in metastatic tumor cells from host organs of another species ^11–16^. Such assays are extremely sensitive and quantitative for xenograft models that transplant human tumor cells onto mice. However, this powerful method is rarely used in syngeneic mouse models. Since the syngeneic tumor cells possess identical genomic DNA as the host, the more-established qPCR methods that leverage species-specific amplification are not applicable. Efforts were also made for the development of qPCR method using cDNA amplification in syngeneic models based on the different expression level of signature genes between metastatic tumor cells and animal host tissues ^15,16^. Yet, the results were more qualitative than quantitative as the signature gene expression of metastatic cells, unlike identical genomic DNA, will be affected by the surrounding microenvironment at different locations ^4,5^.

To address some of these challenges with quantifying spontaneous metastases in syngeneic mouse models, we developed a real time PCR method for quantifying metastatic tumor load in peripheral organs and illustrated the approach using a syngeneic melanoma mouse model to track and quantify spontaneous metastasis after injecting C57BL/6Ncrl or NOD-Scid Gamma (NSG) mice with mouse B16F0 (non-metastatic) cells and B16F10 (metastatic) cells. In order to detect the tumor cells using both bioluminescence imaging and real time qPCR, we introduced modified luciferase gene *Luc2* into B16 cells via lentivirus, designed specific primers for luciferase gene fragment amplification, and developed a real time qPCR method that can quantify tumor cells with luciferase gene insertions. In each qPCR reaction, we can quantitatively detect one Luc2+ tumor cell within 10,000 mouse cells. When we used genomic DNA extracted from the whole mouse lungs, we could quantitatively identify as few as 1.8x10^4^ metastatic cells within the lungs. In comparing bioluminescence imaging and qPCR to detect B16 melanoma metastasis, we found the qPCR method was at least a factor of 10 more sensitive in metastatic cell determination and quantitation. The results from two methods were consistent and supported each other, but qPCR gave a final definitive quantitation for tumor cell metastasis.

## Experimental

### Luciferase-expressing lentivirus packaging

Lentiviral vector pLU-Luc2, expressing a codon-optimized luciferase reporter gene *Luc2*, was kindly provided by Dr. Alexey V. Ivanov (West Virginia University). The *Luc2* gene was originally adapted from Firefly luciferase reporter vector pGL4.10[luc2] (Promega Corp. WI; GenBank accession number: AY738222.1) and moved into the lentiviral vector under the control of a CMV promoter. Standard lentivirus packaging procedure was performed using pLU-Luc2 and two packaging plasmids, psPAX2 (Addgene plasmid #12260) and pCMV-VSG-G (Addgene plasmid #8454), in HEK293T cells. Virus soup was aliquoted and saved at −80°C.

### Cell culture and lentivirus transduction

Mouse melanoma cell lines B16F0 and B16F10 were from American Type Culture Collection (ATCC, Manassas, VA), and maintained in DMEM supplemented with 10% FBS and penicillin/streptomycin at 37 °C, 5% CO2. To create Luc2-expressing mouse melanoma cells, the two cell lines were prepared on 6-well plates, and transduced with lentivirus soup with an estimated MOI (multiplicity of infection) of 5. The virus soup was incubated with cells for 24 hours in the presence of 5.0μg/ml of polybrene (Sigma-Aldrich, St. Louis, MO). The transduced cells were propagated, tested by Western blotting and bioluminescence assay for similar and stable Luc2 expression before injecting into the mice.

### Spontaneous and experimental metastasis assay

Animal experiments described in this study were approved by West Virginia University (WVU) Institutional Animal Care and Use Committee and were performed at the WVU Animal Facility in compliance with the relevant laws and institutional guidelines. 6-week female C57BL/6Ncrl mice were purchased from Charles River Laboratories and 6-week male NOD-Scid Gamma (NSG, Stock No: 005557) mice were from The Jackson Laboratory.

To assay spontaneous metastasis, C57BL/6Ncrl or NSG mice were injected subcutaneously with B16F0-Luc2 (3X10^5^/mouse) or B16F10-Luc2 cells (1.2X10^5^/mouse). For an experimental metastasis assay, NSG mice were injected intravenously with B16F10-Luc2 cells (6X10^4^/mouse). Bioluminescence imaging was performed to quantify tumor burden in vivo or ex vivo at the indicated dates. Animal lungs were then collected and frozen at −80°C for qPCR analysis. Some lungs were fixed with 10% neutral buffered formalin (NBF) (Sigma-Aldrich) as indicated.

### Bioluminescence imaging

For in vivo assay of live animals, mice were injected intra-peritoneally with D-luciferin (Caliper Life Sciences, 150 mg/kg) 10 minutes prior to imaging. For ex vivo assay of animal organs, mice were euthanized 10 minutes after D-luciferin injection. Lungs were dissected and incubated in D-luciferin/PBS solution for imaging. All images were taken using the IVIS Lumina-II Imaging System (PerkinElmer, Waltham, MA) with 1.0 minute capture and medium binning. Living Image-4.0 software was used to process the captured images. Signal intensity was quantified as the sum of all detected photon counts within the region of interest after subtraction of background luminescence.

### Genomic DNA extraction

Unless indicated, genomic DNA was extracted from fresh cultured cells using DNeasy Blood & Tissue Kit (Qiagen, Hilden, Germany) according to manufacturer’s instructions. RNase A was used to ensure RNA-free DNA extraction.

Genomic DNA from mouse lungs was extracted by Proteinase K digestion followed by ethanol precipitation. Briefly, mouse lungs (100-200mg) were ground by a syringe plunger on ice, and digested in 20 times of volume of digestion buffer (10 mM Tris-Cl, pH 8.0, 1mM EDTA, 1.0mg/ml Proteinase K, 0.5% SDS) at 56°C until digesting solution became clear (24-72 hours). RNase A was then added to a final concentration of 100μg/ml. Samples were incubated at 37°C for 30 minutes before extracted with Phenol: Chloroform: Isoamyl Alcohol (25:24:1) twice, and DNA was precipitated by ethanol. DNA pellets were washed with 70% ethanol and resuspended in TE buffer (10 mM Tris-Cl, pH 8.0, 1mM EDTA). DNA concentrations were measured on a NanoDrop-1000 spectrophotometer (Thermo Fisher Scientific, Inc., Waltham, MA) and diluted with Milli-Q water for qPCR analysis.

### Real time quantitative PCR (qPCR) and calculation of Luc2 cell ratio

The qPCR primers for firefly luciferase gene *Luc2* was designed using NCBI Primer-Blast and the primers for mouse prostaglandin E receptor 2 (*Ptger2*) gene is reported previously ^13^. The primer sequences are listed as follows: Luc2 forward, CACCGTCGTA TTCGTGAGCA, Luc2 reverse: AGTCGTACTCGTTGAAGCCG; Ptger2 forward, CCTGCTGCTTATCGTGGCTG, Ptger2 reverse, GCCAGGAGAATGAGGTGGTC.

All real time qPCR assays were performed on a StepOnePlus Real-Time PCR System (Thermo Fisher Scientific) using Power SYBR Green PCR Master Mix (Thermo Fisher Scientific) on a MicroAmp Fast Optical 96-Well Reaction Plate (Thermo Fisher Scientific). Unless otherwise specified, 100ng of genomic DNA was used in a 25μl reaction. Each biological sample was amplified in triplicate, whereby the mean Ct value obtained from the technical replicates was used for final calculation. On each plate, serial dilutions of B16F0-Luc2 genomic DNA were used for *Luc2* and *Ptger2* amplification to create standard curves for the calculation of relative Luc2 DNA and total mouse DNA. The qPCR conditions were as follows: 95°C-10 minutes, 40 cycles of (95°C-30 seconds, 61°C-1minute). The results were analyzed and exported by StepOne software v2.3.

Microsoft Excel 2013 was used to establish gene amplification standard curves (Ct vs. log DNA) for *Luc2* and *Ptger2*. The relative Luc2 DNA amount (Q_Luc2_) and total mouse (Ptger2) DNA amount (Q_mm_) were then calculated as described in the text. The Luc2 cell ratio is calculated as: R = Q_Luc2_ /Q_mm_. R is presented as Luc2 cell number in 10^4^ tissue (lung) cells.

### Calculation of DNA amount, lentivirus insertions, and lung cell numbers

The following formula was used to calculate the molecular weight (in Dalton) of mouse genomic DNA. MW of dsDNA= (number of base pairs) X (650 Daltons / base pair). With a total length of 2671.82 MB in mouse haploid genome (NCBI Mus musculus assembly GRCm38.p5), the weight (in picogram) of mouse genomic DNA per cell (diploid) is calculated as: 2 X 650 X 2.67182 X 10^9^ X 10^12^/ (6.022 X 10^23^) = 5.77 pg. Using this number, we also estimated cell number in an organ. Specifically, the lungs normally weighted about 150mg (C57BL/6Ncrl) or 200mg (NSG) at harvest. An average of 400-600μg of genomic DNA was isolated from the lungs of each animal. Using a value of 500μg for the amount of genomic DNA, we calculated the total cell number in the lungs of each mouse as: 500 X 10^6^/5.77 (pg per cell) = 9X10^7^ cells. Similarly, Perrone et al. reported that lungs from 6-8 week old C57BL/6Ncrl mice contain 1.5X10^8^ cells ^17^. The weight corresponding to a certain number of pLU-Luc2 lentiviral vector molecules was also calculated using a similar approach. As described in the qPCR protocol, serial dilutions of the plasmid were used for *Luc2* amplification to create standard curve and to calculate lentiviral insertion copy numbers.

## Results and discussion

### Specific amplification of firefly luciferase gene *Luc2*

Characterizing spontaneous metastasis of mouse melanoma B16 cells in the syngeneic C57BL/6Ncrl or NSG mice is important as it captures all of the stages associated with tumor cell metastasis in an immunocompetent and immunodeficient context, respectively. Given the genetic similarity between the injected B16 melanoma cells and somatic cells from the host, we expressed the firefly luciferase gene in B16 cells with lentivirus transduction and used bioluminescence imaging to track metastatic B16 in vivo. As it became clear that bioluminescence imaging lacked the sensitivity to reveal micrometastases in distant organs such as lungs and liver, we developed a method to quantify the ratio of the luciferase gene-containing genomic DNA to total mouse genomic DNA in these organs. This ratio represents the relative metastatic cell burden in the assayed organ. The method leverages real-time quantitative PCR to detect the inserted luciferase gene in metastatic cells and a reference mouse gene to quantify total mouse DNA.

NCBI Primer-Blast was used to design oligonucleotide sequences for qPCR of *Luc2* gene. After testing several primer pairs for efficiency and specificity, we eventually selected a pair that amplifies a 173-base-pair DNA fragment from Luc2-containing cells, with no other nonspecific DNA fragment generated from regular mouse or tissue genomic DNA (Fig. 1). For quantifying mouse DNA, we selected a primer pair that amplifies a 189-base-pair DNA fragment in mouse prostaglandin E receptor 2 (*Ptger2*) gene in a genetically stable region as reported (Fig. 1) ^13^. To extend the approach to xenograph mouse models, amplifying this DNA fragment with species-specific primer pairs could be used to distinguish between human and mouse genomic DNA and quantitatively determine the relative ratios of the mixture ^13^.

**Fig. 1.**
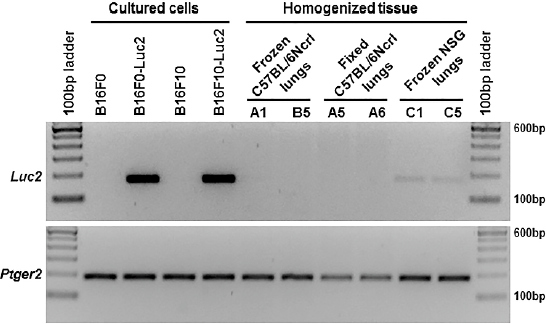
Specific amplification of *Luc2* and *Ptger2* gene fragments with genomic DNA from mouse cells and lungs. 100 ng of genomic DNA was used as template in each 25μl reaction and PCR was performed for 30 cycles with primer pairs as described in Experimental section. The products were resolved on 2.5% agarose gel. The labels for the lungs of C57BL/6Ncrl and NSG mice represent individual animals described in Table 2, in which Luc2 positive cells were detected in the lungs of mouse C1 and C5 using bioluminescence imaging.

### Generation of standard curves and calculation of Luc2 DNA

The real-time qPCR assay was designed to estimate the ratio of Luc2 DNA contained within total mouse DNA, representing the percentage of metastatic B16-Luc2 cells within a total population of mouse cells. We used a single-plex PCR format given the potential wide dynamic range in target concentrations. To relate the amount of genomic DNA with Ct values, we generated two standard curves for both amplicons using a defined amount of luciferase-containing DNA and a defined amount of mouse reference DNA. Using these standard curves, an amount of Luc2^+^ DNA and an amount of mouse DNA (e.g. Ptger2^+^ DNA) from the Ct values were obtained for uncharacterized genomic DNA samples. The ratio of the two amounts corresponds to the metastatic cell burden in assayed cell mixture or organs.

Serial DNA dilutions obtained from cultured cells were used as a qPCR template to investigate the assay sensitivity, reliability and linearity (Table 1 and Supplemental Fig. S1 and S2). The qPCR plots and melting curves for *Luc2* gene showed specific and quantitative amplification from B16F0-Luc2 genomic DNA, whether the DNA were serially diluted with water (Samples T2-T6 and Supplemental Fig. S1A and S1B), or serially diluted with B16F0 genomic DNA (Sample U2-U6 and Supplemental Fig. S1E and S1F), demonstrating the specificity of the amplification. There was also no amplification for *Luc2* gene when B16F0 genomic DNA was used as template alone (Sample S1-S7 and Supplemental Fig. S1C and S1D). When 1.0μg DNA was used as template in the 25 μl reaction, the Ct value was higher than that when 0.1μg DNA was used (19.94 in Sample T1 versus 19.36 in Sample T2), suggesting that the qPCR reaction was partially inhibited by the excess amount of DNA. In the most dilute sample (Sample T7), 1.0 pg DNA resulted in inconsistent amplification or no amplification in duplicates, indicating the sensitivity limit of the assay. Similarly, the qPCR plots and melting curves for *Ptger2* gene also showed specific and quantitative amplification from either B16F0-Luc2 genomic DNA (Samples T2-T6 and Supplemental Fig. S2A and S2D), or B16F0 genomic DNA (Samples S2-S6 and Supplemental Fig. S2B), or mixed genomic DNA (Samples U2-U6 and Supplemental Fig. S2C). In amplifying *Ptger2*, we also observed similar upper and lower limits for DNA template abundance as observed with *Luc2* amplifcation (1.0μg and 1.0pg, respectively).

**Table 1.** Generation of standard curves for real-time quantitative PCR to calculate Luc2^+^ DNA and total mouse (Ptger2) DNA.

| Sample | B16F0-*Luc2* DNA (pg) | B16F0 DNA (pg) | Ct for *Luc2* | Ct for *Ptger2* |
| --- | --- | --- | --- | --- |
| T1 | 1,000,000 | - | 19.94 | 20.84 |
| T2 | 100,000 | - | 19.36 | 20.95 |
| T3 | 10,000 | - | 22.66 | 24.02 |
| T4 | 1,000 | - | 25.98 | 27.29 |
| T5 | 100 | - | 29.44 | 30.64 |
| T6 | 10 | - | 33.29 | 33.14 |
| T7 | 1 | - | >34 or n/d | >34 or n/d |
| S1 | - | 1,000,000 | n/d | 20.76 |
| S2 | - | 100,000 | " | 21.02 |
| S3 | - | 10,000 | " | 24.19 |
| S4 | - | 1,000 | " | 27.70 |
| S5 | - | 100 | " | 30.86 |
| S6 | - | 10 | " | 33.31 |
| S7 | - | 1 | " | >34 or n/d |
| U2 | 100,000 | - | 19.32 | 20.94 |
| U3 | 10,000 | 90,000 | 22.91 | 20.96 |
| U4 | 1,000 | 99,000 | 26.04 | 20.96 |
| U5 | 100 | 99900 | 29.41 | 21.21 |
| U6 | 10 | 99990 | 33.29 | 21.09 |

Almost overlapping results were achieved for *Luc2* amplification using B16F0-Luc2 DNA serially diluted with either water or B16F0 genomic DNA (Table 1 and Fig. 2A, sample T2-T6 and U2-U6). With these two datasets, we plotted the mean Ct value for *Luc2* amplifications against the log of the DNA template abundance, and created a linear relation between 0.1μg to 10pg of input of B16F0-Luc2 genomic DNA, with a R^2^ value of 0.9989 (Fig. 2A). Outside this range, we may still detect *Luc2* gene (Sample T1 and T7), but the Ct values can’t be used for quantitative interpretation. For instance, less than 10 pg of input increased the variability among technical replicates, which is consistent with stochastic effects introduced by low template numbers. Using the standard curve, we inverted the relationship to calculate the amount of Luc2 DNA for a given Ct value within the linear range. For example, the curve in Fig. 2A, y =−3.4531x + 36.53, was switched to x = (36.53−y) / 3.4531 (y is the mean Ct, x is log DNA). The Luc2 DNA was calculated as Q_Luc2_ = 10^(36.53− y)^ / 3.4531 (Q_Luc2_ is Luc2 DNA amount, in picogram). Similar linear standard curve was established for *Ptger2* amplification using B16F0-Luc2 (Samples T2-T6) and B16F0 (Samples S2-S6) genomic DNA for the calculation of relative mouse DNA (Qmm), with DNA input between 0.1μg to 10pg for quantitation purpose (Fig. 2B). Collectively, the linear regression results suggest that a log-linear relationship exists between template abundance and Ct values for both amplicons and that the results are highly reproducible.

**Fig. 2.**
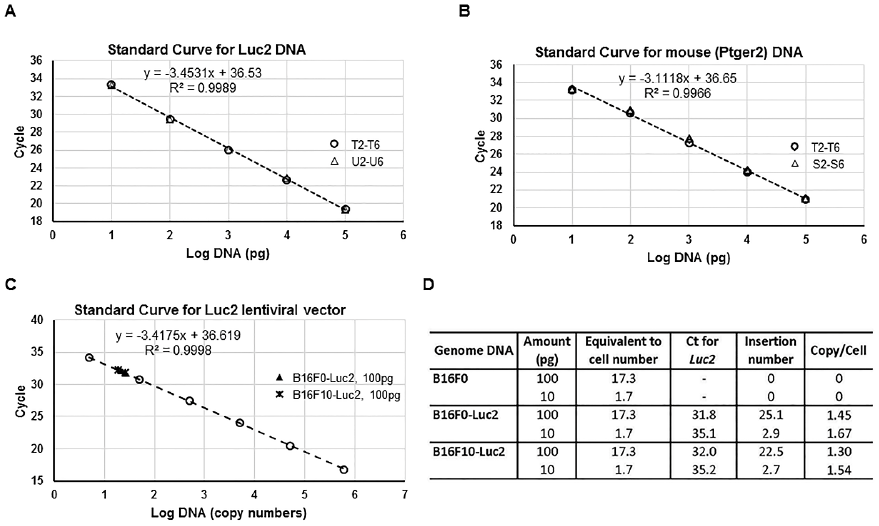
Standard curves with different genomic DNA samples show the linearity and consistence of real-time quantitative PCR for both *Luc2* and *Ptger2* genes. (A) The mean Ct (y-axis) of the serial Luc2+ genomic DNA dilutions in Table 1 (Group T and Group U, without or with addition of Luc2^−^ DNA) were plotted against log values (x-axis) of DNA input (in picogram) to create a Luc2 DNA standard curve. The data points from two groups were indicated: ○, group T; Δ, group U. (B) The mean Ct (y-axis) of the serial Luc2^+^ and Luc2^−^ genomic DNA dilutions in Table 1 (Group T and Group S) were plotted against log values (x-axis) of DNA input (in picogram) to create a Ptger2 DNA standard curve. The data points from two groups were indicated: ∘, group T; Δ, group S. (C) The mean Ct (y-axis) of the serial pLU-Luc2 lentiviral plasmid dilutions were plotted against log values (x-axis) of DNA input (in plasmid copy numbers) to create a linear standard curve for Luc2 lentiviral vector. The data points from sample groups were indicated: ∘, lentiviral plasmid; ▲, 100pg of B16F0-Luc2 genomic DNA; *, 100pg of B16F10- Luc2 genomic DNA. (D) Calculation of lentiviral inserted copy numbers from the Ct values using indicated amount of DNA from B16F0, B16F0-Luc2 or B16F10-Luc2. The calculated DNA amount per mouse cell (diploid) is 5.77pg (see calculation in Experimental section), hence the cell number for 100pg of DNA is: 100/5.77=17.3.

To estimate the degree of *Luc2* insertion into the genome, we used serial dilutions of the original lentiviral vector to create a similar standard curve for lentiviral copy numbers (Fig. 2C). Based on the genomic DNA input, we calculated the lentiviral copy number in B16F0-Luc2 and B16F10-Luc2 cells (Fig. 2D). The insertion number for both is about 1.5 per cell (2 haploid genomes), suggesting our lentiviral transduction efficiency was consistent. Given the similar Luc2 insertion number, we also used the standard curve created from B16F0-Luc2 DNA to calculate the amount of B16F10-Luc2 DNA for assaying spontaneous metastasis of B16F10-Luc2 cells.

Using the three standard curves, we estimated the sensitivity of this qPCR assay to quantify the metastatic cell load in a particular tissue. By operating in the linear range of template abundance, we can detect Luc2 DNA if present in as low as 10 pg of Luc2+ tumor cell DNA out of the maximal 100 ng of total mouse tissue DNA as input. Using 5.77 pg of DNA per cell, this corresponds to 1.7 Luc2+ tumor cells, which collectively contain 2.6 copies of Luc2 lentiviral insertions, and 17,000 tissue cells. Thus, the sensitivity of the assay for detecting Luc2+ tumor cells present within a homogenized tissue is 1 metastatic cell out of 10,000 cells. We also note, however, that storing conditions prior to extracting the genomic DNA from the tissue samples can degrade the overall DNA quality such that the detection limit for frozen lungs decreases to 1 in 5,000 cells and for formalin-fixed lungs decreases further to 1 in 100 cells (data not shown). We estimated that, from mice at the end point of our experiments, each lung contained about 9X10^7^ cells (see Experimental for calculation), this would define our detection limit for Luc2 cells in mouse lungs as 1.8X10^4^ for frozen samples, and 9X10^5^ for formalin-fixed samples.

The sensitivity of this qPCR assay has the potential to be improved. Our standard genomic DNA input in the assay was 100ng, linearity seemed to be degraded at higher DNA input amounts. Optimizing PCR reaction conditions further to increase DNA input while still keeping a linear response over the dynamic range will possibly increase the assay resolution by several folds. Another practical way to increase assay resolution is to increase lentiviral insertion copy numbers by using higher titers of virus and by performing multiple rounds of transductions. However, increasing the lentiviral insertion numbers runs the risk of changing the tumor cell genome and the resulting cellular phenotype with damage from insertions.

### Detection of spontaneous lung metastasis with bioluminescence imaging and qPCR

Following development of the assay and sensitivity metrics, bioluminescence imaging and qPCR were used to evaluate spontaneous lung metastasis in two different scenarios. First, we compared spontaneous lung metastasis in C57BL/6Ncrl mice that received subcutaneous injection of B16F0-Luc2 (Table 2, Experimental Group A) or B16F10-Luc2 cells (Table 2, Exp. Group B). In vivo selection was used to develop these variants of the parental B16 cell line where B16F10 cells exhibit a high metastatic potential while B16F0 cell do not ^18^. The second scenario was to compare spontaneous lung metastasis following subcutaneous injection of B16F0-Luc2 cells in C57BL/6Ncrl mice (Exp. Group A), which are immunocompetent, and in NSG mice (Table 2, Exp. Group C), which are severely immunocompromised.

**Table 2.** Summary of qPCR detection for melanoma spontaneous and experimental lung metastasis in four groups of mice.

| Exp Group | Mouse Strain | Injection Method | Tumor Cell | Injected Number | Mouse ID | Harvest Date (DPI*) | Luc2 Cells in 10<sup>4</sup> Lung Cells |
| --- | --- | --- | --- | --- | --- | --- | --- |
| A | C57BL/6Nrl | Subcutaneous | B16F0-Luc2 | $3 \times 10^5$ | A1 | 21 | < 1.0** |
|  |  |  |  |  | A2 | 23 | < 1.0** |
|  |  |  |  |  | Other 13 mice | 15-25 | 0 |
| B | C57BL/6Nrl | Subcutaneous | B16F10-Luc2 | $1.2 \times 10^5$ | B1 | 22 | 24 |
|  |  |  |  |  | B2 | 22 | 10 |
|  |  |  |  |  | B3 | 22 | 3.1 |
|  |  |  |  |  | B4 | 22 | 3.7 |
|  |  |  |  |  | B5 | 22 | 0 |
|  |  |  |  |  | B6 | 22 | 0 |
| C | NSG | Subcutaneous | B16F0-Luc2 | $3 \times 10^5$ | C1 | 21 | 27.8 |
|  |  |  |  |  | C2 | 21 | 1.4 |
|  |  |  |  |  | C3 | 20 | 1.8 |
|  |  |  |  |  | C4 | 21 | 5.7 |
|  |  |  |  |  | C5 | 21 | 10.4 |
| D | NSG | Intravenous | B16F10-Luc2 | $6 \times 10^4$ | D1 | 16 | 193.3 |
|  |  |  |  |  | D2 | 16 | 391.7 |
|  |  |  |  |  | D3 | 16 | 363.8 |
|  |  |  |  |  | D4 | 16 | 276.5 |

\*DPI: Days Post Inoculation; \*\*<1.0: Luc2 DNA was detected in only one or two of the three replicates, the $C_t$ values of the positive amplifications were higher beyond the linear standard curve for Luc2 DNA.

In comparing C57BL/6Ncrl mice receiving either B16F0-Luc2 or B16F10-Luc2 cells, we could not detect any spontaneous lung metastasis in vivo with bioluminescence imaging at any point following subcutaneous injection of 3x10^5^ B16F0-Luc2 cells, despite shielding of bioluminescence derived from large primary tumors (data not shown, but see Fig. 4A, mouse B4). Ex vivo bioluminescence imaging and qPCR were then used as end point assays to assess melanoma metastasis in lungs. In mice from Exp. Group A, spontaneous lung metastasis were not detected with ex vivo bioluminescence lung imaging (Fig. 3A), but we did detect Luc2 cells with qPCR in lungs of 2 out of 15 mice in the group. While the melting curves suggested a single amplicon, the Ct values from qPCR were greater than 34, variable among technical replicates, and, therefore, outside the range for quantitation (Table 2, mouse A1 and A2). Such low metastatic ratio in mice (13.3%, 2 out of 15) and low Luc2 cell ratios in lungs with metastasis (i.e., detectable but less than 1 cell in 10^4^) were consistent with the nature of B16F0 as a non-metastatic melanoma cell line ^18^. In contrast, subcutaneous injection of fewer B16F10-Luc2 cells (1.2x10^5^ cells) resulted in detecting spontaneous lung metastasis with ex vivo bioluminescence imaging in 1 out of 6 samples in group B (Fig. 3B). At the experimental endpoint, qPCR detected spontaneous lung metastasis in 4 out of the 6 mice, about 66.7% of metastatic ratio in this group (Table 2). The fact that the sample B1, the only lungs with metastasis revealed by bioluminescence imaging, possessed the highest lung metastatic cell ratio (24 in 10^4^ lung cells) illustrates consistency between the two assays. The results from these two groups also suggested that the qPCR assay was at least a factor of 10 more sensitive and more quantitative than ex vivo bioluminescence imaging.

**Fig. 3.**
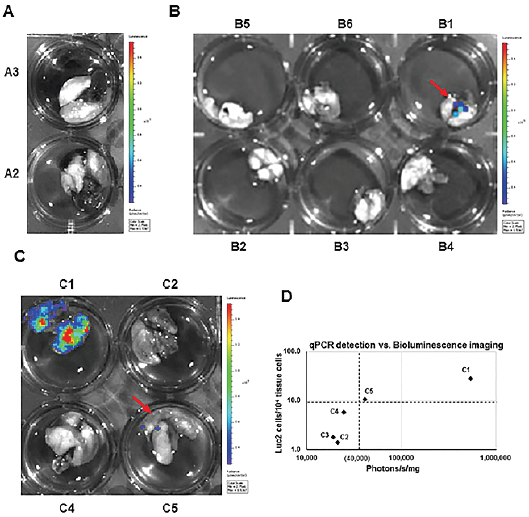
Detection of spontaneous lung metastasis with bioluminescence imaging. C57BL/6Ncrl or NSG mice with subcutaneous injection of B16F0-Luc2 or B16F10-Luc2 cells were terminated for ex vivo bioluminescence imaging at indicated time as listed in Table 2. Representative results of ex vivo bioluminescence imaging of lungs obtained from mice in group A (panel A), in group B (panel B), and in group C (panel C). (D) Correlation of qPCR detection with bioluminescence imaging sensitivity. The whole lung surface photon counts (photons/s) in each of group C mice were measured and normalized to lung weight (in mini gram), the resulted number was used as x- coordinate of the sample. With metastatic cell ratio (in 10^4^ lung cells) as y-coordinate, each sample was plotted in the diagram.

As part of a broader effort to evaluate the role of mouse immune response against melanoma metastasis, we also subcutaneously injected 3x10^5^ B16F0-Luc2 cells into immunocompromised NSG mice (Exp. Group C), which we compared against the response using C57BL/6Ncrl mice in Exp. Group A (Table 2 and Fig. 3C). Interestingly as it was different from the results for group A, we saw spontaneous lung metastasis with ex vivo bioluminescence imaging in 2 out of 4 samples, one of them (C1) gave out very strong signals (Fig. 3C). qPCR revealed that all 5 out of 5 mice possessed lung metastases, and C1 had the highest lung metastatic cell ratio (27.8 in 10^4^ lung cells). Interestingly, there was a much higher incidence (i.e. 100%) of spontaneous lung metastasis in NSG mice relative to C57BL/6Ncrl mice, which suggests that the mouse immune response represses melanoma metastasis. In relating qPCR-determined Luc2 tumor cell ratio with an estimate of metastatic tumor load using bioluminescence imaging (Fig. 3D), we observed a threshold for detecting metastases by bioluminescence imaging at 10 metastatic cells in 10^4^ lung cells, which corresponded to 40,000 photons/s/mg for bioluminescence imaging. Above the threshold, both methods could detect metastases, while below the threshold, only qPCR is sensitive enough to detect metastases. In addition, these results also supported our previous conclusion that the qPCR assay has much higher sensitivity and resolution than bioluminescence imaging in detecting spontaneous metastasis.

### Comparison of spontaneous and experimental lung metastasis detection

Experimental metastasis assays are more commonly used partially due to difficulty for spontaneous metastatic tumor cell detection ^6,7^. To further reveal the potential of qPCR in metastatic tumor cell detection, we performed an experimental metastasis assay in a group of NSG mice using intravenous injection of 6x10^4^ B16F10-Luc2 cells in the tail vein (Table 2, Exp. group D). In the experimental, both bioluminescence imaging and qPCR were used for the mice, and results were compared with C57BL/6Ncrl mice receiving a subcutaneous injection of B16F10-Luc2 cells (Exp. Group B). A comparison of the imaging results for representative samples are shown in Figure 4.

**Fig. 4.**
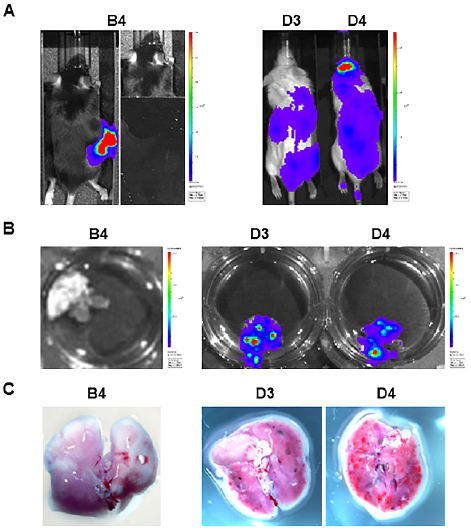
Comparison of spontaneous and experimental lung metastasis detection with bioluminescence imaging and close-up photography. Summaries of representative mice for spontaneous metastasis (B4) and experimental metastasis (D3 and D4) were listed in Table 2. (A) In vivo bioluminescence imaging one day before the mice were terminated as listed in Table 2. On the right side of panel B4, the primary melanoma was shielded for better detection of lung metastases. (B) Ex vivo bioluminescence lung imaging. (C) Photos for black B16F10-Luc2 macrometastases on lung surfaces.

After tail vein injection, B16F10-Luc2 cells spread through bodies and grew rapidly, especially in locations such as brain, lungs, liver, kidney and intestines (Fig. 4A). Due to the high tumor burden in the experimental metastasis assay, the experiment ended at day 16 compared to the regular 3 weeks for spontaneous metastasis assays.

While the spontaneous metastatic cells could not be detected with bioluminescence imaging in any of the mice from Exp. Group B, strong bioluminescence signals were observed for mice in Exp. group D associated with the lungs both in vivo and ex vivo (Fig. 4A and 4B). When the lungs were dissected, visible black macrometastases were observed around surface of lungs (Fig. 4C, compare B4 with D3 and D4), though they were still much smaller than the regular lung metastatic nodules. qPCR was performed using DNA obtained from the lungs (Table 2). The calculated lung metastatic cell ratio for those group D mice was about 200-400 B16F10-Luc2 cells per 10^4^ lung cells (i.e., 2-4% of total lung cells).

Collectively, these in vivo studies provided a clear biological context for evaluating and quantifying spontaneous and experimental metastasis using different detection methods. While metastatic cells need to take 2-4% of total lung cells to form visible macrometastases (Fig. 4), an estimated metastatic cell burden of more than 10% in lungs is needed to form the regular metastatic tumor nodules for manual counting (data not shown). Bioluminescence imaging greatly increased detection sensitivity, as it only needs about 0.1-0.2% of metastatic cells in lungs for detection ex vivo (Fig. 3). As a further improvement in sensitivity, the qPCR assay developed in this work can quantitatively detect as low as 0.01% of metastatic cells in lungs, and qualitatively detect even fewer cells (Table 2).

## Conclusions

In this work, we developed a real time quantitative PCR method to detect spontaneous metastasis in the syngeneic mouse melanoma model. To track and quantify spontaneous metastatic B16 mouse melanoma cells in mice, we introduced firefly luciferase gene *Luc2* into B16 cells with lentivirus. In addition to standard method of bioluminescence imaging to detect luciferase-expressing metastatic cells, we designed specific primers for *Luc2* gene amplification, and developed a real time qPCR method that can quantify *Luc2* positive genomic DNA with lentiviral insertions. An internal control gene (*Ptger2*) was used to normalize the quality of genomic DNA, and the ratio of the Luc2 positive cells within total mouse cells was then determined.

In vivo assays of spontaneous mouse metastasis showed that the qPCR method was highly sensitive and able to detect a very small amount of metastatic cells in mouse lungs that could not be revealed with regular bioluminescence imaging. We estimated that the qPCR method was at least one order of magnitude more sensitive in quantifying metastatic cell burden than bioluminescence imaging. While the results from two methods were consistent, qPCR gave a final definitive quantitation for tumor cell metastasis. We highly recommend this qPCR assay as a standard procedure together with bioluminescence imaging as a high-resolution, end-point method for final metastatic cell quantitation. Given the rapid nature of tumor growth using the B16 model, assays with improved sensitivity can provide better insight into biological mechanisms that underpin tumor metastasis.

## Acknowledgements

This work was supported by National Science Foundation (NSF CBET-1644932 to D. J. K.) and National Cancer Institute (NCI 1R01CA193473 to D. J. K.). The content is solely the responsibility of the authors and does not necessarily represent the official views of the NSF or NCI. The authors thank Alexey V. Ivanov from West Virginia University for providing luciferase-expressing lentiviral vector pLU-Luc2. Small animal imaging and image analysis were performed in the West Virginia University Animal Models & Imaging Facility, which has been supported by the West Virginia University Cancer Institute and NIH grants P20 RR016440, P30 GM103488 and S10 RR026378.

